# A hybrid micro-ECoG for functionally targeted multi-site and multi-scale investigation

**DOI:** 10.1101/2024.12.18.629147

**Authors:** Patrick Jendritza, Rickard Liljemalm, Thomas Stieglitz, Pascal Fries, Christopher Murphy Lewis

## Abstract

Brain function relies on coordinated activity across a wide range of spatial and temporal scales. The activity of single neurons depends on their unique pattern of local and long-range connectivity and thus on large-scale patterns of distributed activity across brain-wide networks. Understanding integrated, whole brain function requires new tools capable of recording from anatomically connected populations in distributed brain areas to bridge local and global dynamics. Here, we present high-density, micro-electrocorticography arrays that facilitate multi-scale studies of brain activity. The hybrid arrays integrate the desirable features of silicone elastomers and polyimide films. The silicone elastomer superstructure provides optical transparency and permits repeated penetration with rigid intracortical arrays. The polyimide films enable fine feature definition through photolithography. This combination facilitates high-throughput functional mapping of brain areas to define functionally characterized targets and permits targeted insertion of depth electrodes through the same array for dense local sampling. We demonstrate their suitability for functional mapping of cortical regions in rats, cats and marmosets and the benefit of the resulting functional maps for targeting functionally defined populations for multi-area laminar recordings. Finally, we demonstrate the utility of the hybrid µECoG for local optogenetic stimulation and its use in investigating cortico-cortical interactions through feedforward optogenetic stimulation. Together, these capabilities make the hybrid µECoG a compelling tool for systems neuroscience.

## Introduction

The introduction of new optical and electrophysiological tools, as well as the widespread adoption of functional brain imaging, has prompted a paradigm shift in systems neuroscience away from a focus on individual neurons and brain areas and towards a greater appreciation of coordinated activity across brain-wide networks.^[1–10]^ While classic studies focused on the functional characterization of single neurons and brain areas,^[11–14]^ advances over the past decades have demonstrated the importance of understanding the coordination of neuronal populations within local microcircuits and the integration of activity across distributed, brain-wide networks.^[15–19]^ Indeed, while neurons are powerful computational units,^[20–22]^ the brain’s remarkable capabilities are achieved through communication within and between neuronal groups, and the functional characteristics of single neurons can be better understood in light of local and global activity patterns.^[18,23–25]^ These insights have been critically aided by methodological advances that enable multi-scale and multi-modal measurement and perturbation of brain activity^[7,15,16,26–28]^ to map neuronal activity both densely within a local microcircuit and broadly, across relevant brain regions. However, to date, our understanding of distributed processing in the brain is fundamentally limited by our ability to record simultaneously from multiple, anatomically connected neuronal populations across brain-wide networks with high spatial and temporal precision.

Currently, simultaneous inter-areal recordings of communicating neurons require the laborious search for functionally paired locations in areas of interest guided by anatomy or based on functional maps determined using imaging in different experimental sessions. The difficulty of localizing the specific subpopulations of anatomically connected, communicating neurons, especially in large brains, has hampered the investigation of inter-areal communication, particularly at the cellular scale.^[29,30]^ New tools and techniques are required to investigate the principles of multi-scale brain activity that bridge cellular activity within local circuits to distributed patterns of activity across brain-wide networks.^[31,32]^ The rapid development of optical methods to functionally image populations of neurons^[7,33,34]^ in combination with viral and genetic methods to target specific neuron classes based on their cell-type or their pattern of inter-areal axonal projections^[35–37]^ have made it possible to monitor specific projection neurons and functionally characterize the axons or somata of projecting cells in a directed manner.^[38–40]^ However the field-of-view of most microscopes is still limited to imaging one local population (but see: ^[41–43]^), and population imaging does not yet permit the reliable recording of single action potentials at the millisecond temporal scale. In parallel, advances in the design and fabrication of intracortical multielectrode arrays have greatly increased the number and density of recording sites, permitting high-yield recordings of neuronal activity within areas of interest.^[44,45]^ However, it is difficult to scale the placement of multiple multielectrode arrays to cover brain-wide networks for simultaneous recording (but see: ^[46,47]^). In contrast, micro-electrocorticography (µECoG) provides a means to record from distributed brain regions with broad coverage and high spatial and temporal resolution, but it is limited to signals at the surface of the brain.^[48–50]^ The complementary features of these two approaches offer a compelling combination: functional maps derived from the large-scale, high-resolution mapping afforded by µECoG can be used to specify candidate targets for dense local sampling across the cortical depth with intracortical multi-electrode arrays^[51,52]^. Functional mapping and targeted recording can be combined across brain areas, where functional topography partially determines the likelihood of communicating subpopulations, thereby increasing the probability of recording from anatomically connected neuronal populations and aiding the investigation of inter-areal communication.^[53,54]^ This is particularly the case for areas with clear functional topography, such as retinotopically organized areas of the visual cortex^[55]^, tonotopically organized areas of the auditory cortex^[56]^, or somatotopically organized regions of the somatosensory and motor cortex.^[57,58]^

We present a hybrid µECoG for performing large-scale, high-resolution functional characterization of targeted brain areas in combination with dense, functionally targeted local sampling from regions of interest. The combination of high-resolution multi-electrode arrays fabricated on polyimide thin-films with transparent, flexible and elastic silicone elastomers (Polydimethylsiloxane, PDMS) enables high-resolution electrical sampling from the cortical surface, while preserving optical access to the underlying cortex. Further, the flexibility, elasticity and self-sealing nature of PDMS permits repeated penetration with rigid linear arrays to perform recordings of neuronal activity across the cortical depth in functionally identified populations.^[59,60]^ The hybrid µECoGs are conformal^[61]^ to the delicate cortical surface and provide dense functional information spanning large regions of interest at the scale of cortical areas (tens to hundreds of square millimeters). We demonstrate the utility of the design in three different species (rat, cat, and marmoset) and show that the arrays are capable of recording both the low frequency, local field potential, as well as the aggregate spiking activity of superficial neurons recorded from the cortical surface.^[50]^ These signals can be used to functionally characterize the local population across the cortical surface, enabling the targeting of specific populations for depth recordings with multielectrode arrays inserted into the brain while the µECoG remains in place. We further show that the optical transparency of the hybrid µECoG permits local and inter-areal activation using optogenetics.^[62]^ This approach enables the investigation of local circuit activity in relation to ongoing, large-scale, multi-area brain states, as well as interference with functionally characterized populations and optogenetic activation of inter-areal projection neurons. Such tools promise to open new avenues for investigating the integration of cellular activity with whole-brain activity patterns.

## Results

### A hybrid µECoG for functionally targeted multi-scale investigation

We developed a hybrid µECoG array to enable high-resolution functional characterization of brain areas from the cortical surface, while permitting combined recordings with depth electrodes and optical access for optogenetic perturbations or imaging. High resolution is realized by flexible linear electrode arrays fabricated through photolithography on polyimide thin films, while optical and depth access are provided by a superstructure formed from silicone elastomer (Figure 1a). Silicone elastomers provide a few other practical advantages over a pure thin-film ECoG array. Silicone is soft and conformal, gently accommodating to the cortical surface, avoiding edema and sealing the exposed cortex, while avoiding wrinkling or buckling which would be difficult or impossible to achieve with the polyimide alone. Once fabricated, the polyimide linear arrays can be easily and flexibly assembled to match the size of specific brain areas or the brains of different species. To illustrate the flexibility and utility of these arrays, we assembled different configurations and used them to map the retinotopy of visual cortical areas and investigate local and inter-areal interactions across the visual cortical hierarchy of three model systems: the rat, the marmoset and the cat (Figure 1b). The arrays can be configured to have higher or lower recording density depending on needs and can be flexibly positioned to record from brain areas of interest. Once in place on the cortical surface, the hybrid µECoG allows repeated penetration with depth electrodes and provides optical access for simultaneous optogenetic stimulation, or imaging (Figure 1c). To target populations of interest, or to increase the chances of recording subpopulations of anatomically connected neurons, functional maps can be estimated from the µECoG recordings, and sites of interest with similar tuning or response properties can be selected for simultaneous laminar recording with penetrating linear arrays (Figure 1d). We introduce the fabrication of hybrid µECoG arrays and illustrate their utility for investigating local and inter-areal cortical interactions in the visual system of anesthetized rats, cats, and marmosets.

**Figure 1:**
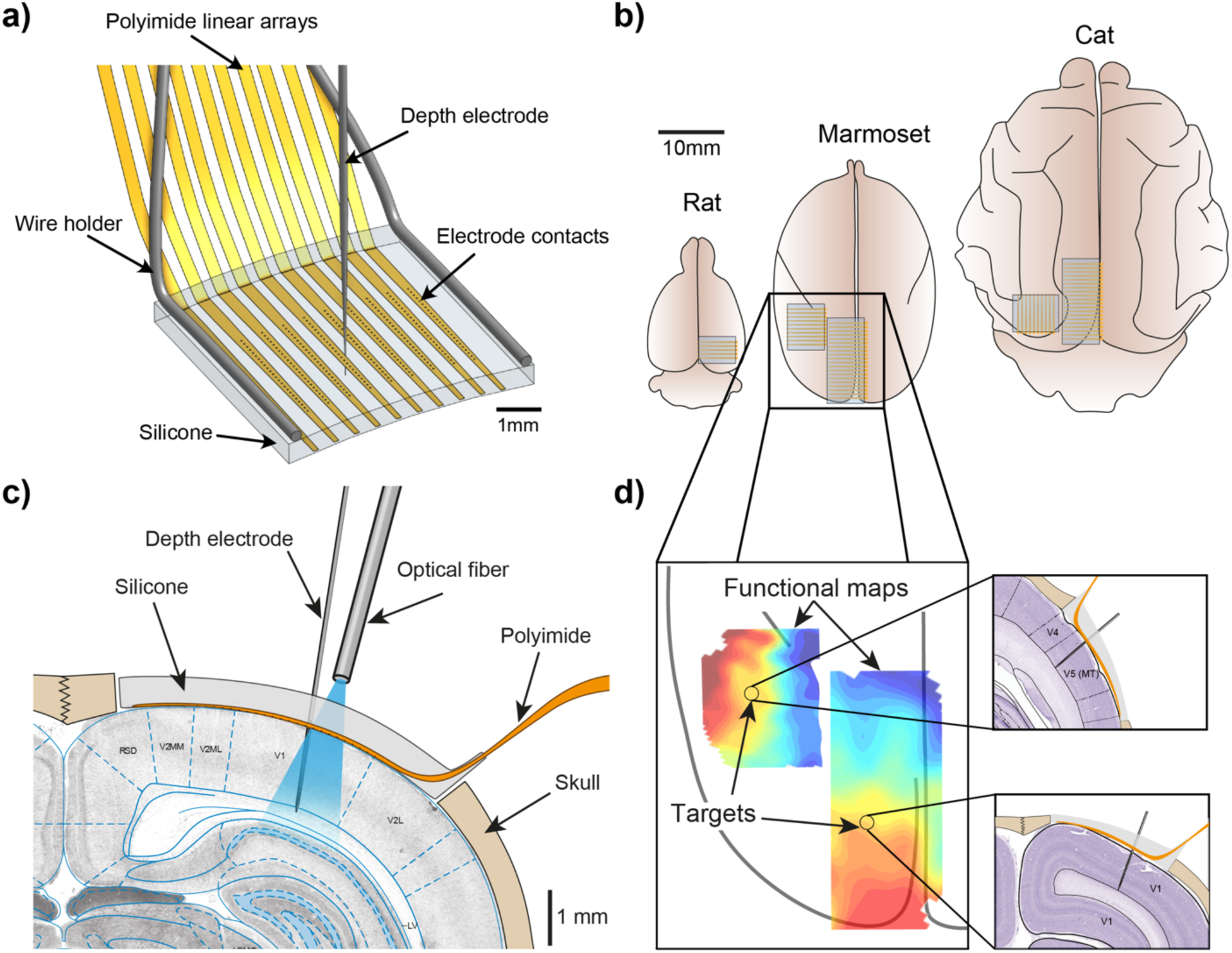
A hybrid µECoG for functionally targeted, multi-scale investigation. **a)** Illustration of a 240-channel hybrid µECoG array showing ten 24-electrode polyimide thin-film linear electrode arrays regularly arranged and encapsulated in a silicone elastomer and secured to a wire holder for placement on the brain. A depth electrode is depicted penetrating the hybrid µECoG at a desired location. **b)** Illustration of hybrid µECoG arrays placed on the rat, marmoset and cat brain to indicate the scale of recordings and coverage of different array designs on the model organisms used in this study. **c)** Schematic of a hybrid µECoG in place on the rat brain with a penetrating linear depth array and an optical fiber to perform optogenetic activation of local neurons or projections. **d)** Schematic illustration of functional maps derived from µECoG recordings from the cortical surface and target locations picked by similarity of functional maps. (Right) Illustration of depth linear electrode arrays inserted through the µECoG to record laminarly resolved neuronal activity from the target locations in the marmoset brain.

### Fabrication of a hybrid µECoG

We fabricated µECoG arrays in a two-step process, using two biocompatible polymers to produce a hybrid array that combines the complementary advantages of each. First, Micro-Electro Mechanical Systems (MEMS) processes were used to fabricate flexible high-density linear electrode arrays on polyimide thin films (Fig 2A).^[48,63–65]^ MEMS fabrication with polyimide permits small feature sizes, flexibility and biocompatibility, but polyimide is not transparent, has some autofluorescence, and cannot be penetrated by sharp electrodes. To achieve the desired features, these high-density polyimide arrays were integrated into a silicone elastomer (Polydimethylsiloxane, PDMS) that can act as a dural replacement, is optically transparent, stretchable and flexible and can be repeatedly penetrated by sharp electrodes (Fig 2B).^[51,66,67]^ However, in contrast to polyimide, it is challenging to achieve high-resolution, small features on PDMS using standard MEMS fabrication processes, because of its elasticity and poor adhesion with photoresist. In addition, PDMS is not compatible with standard MEMS cleanroom manufacturing since residues contaminate machines and surfaces and seriously deteriorate process performance. Using this two-step process also permitted us to decouple the precision lithography necessary to fabricate high-density arrays, from the modular assembly of multiple thin-film arrays into the final hybrid µECoG. By using this two-step process, a single thin-film design can be recombined into multiple µECoG arrangements that can be assembled in the lab, without the need for further work in the clean room. The µECoG can thus be flexibly assembled to match the geometry of the brain area and species of choice and reassembled for different experiments, eliminating the need for costly photolithography mask redesigns and other time-intensive cleanroom procedures.

**Figure 2:**
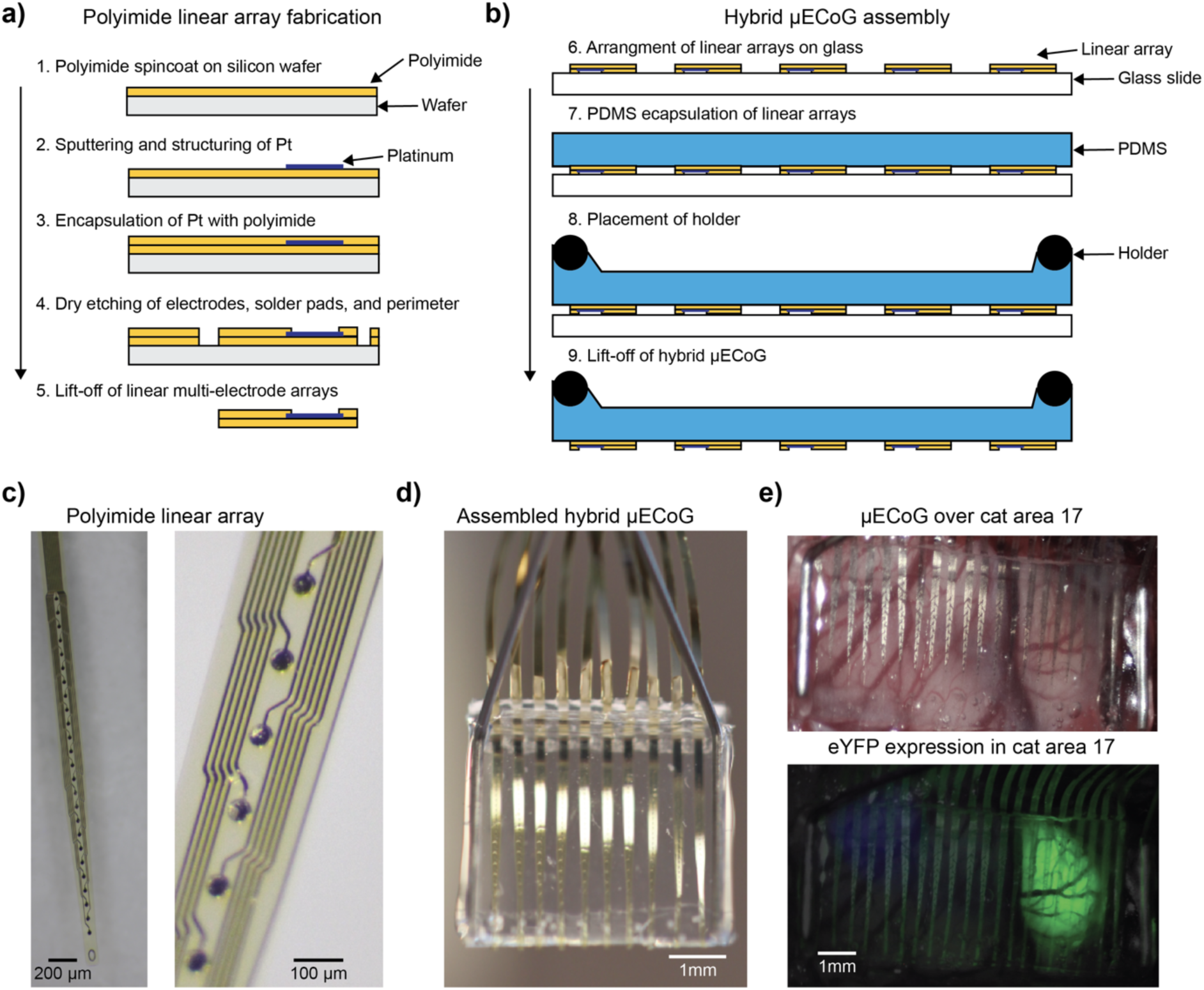
Fabrication of the hybrid µECoG. **a)** Illustration of the fabrication sequence for the polyimide thin-film electrode arrays. **b)** Illustration of the assembly of the hybrid µECoG by regular arrangement of polyimide thin-film arrays and encapsulation with silicone (PDMS) and attachment of the wire-holder. **c)** Photomicrographs of a 24-channel polyimide linear array with 35 µm diameter platinum electrode sites. **d)** Photograph of an assembled hybrid µEcoG with ten 24-channel polyimide thin-film arrays. **e)** Photograph of a hybrid µECoG with twenty 12-channel polyimide thin-film arrays placed over the cat primary visual cortex (top), and epifluorescence image (bottom) showing the virally expressed fluorescence protein (eYFP).

In this process, the design of the high-resolution thin-film electrode arrays is the first step, and it is desirable to choose a layout that can be flexibly integrated into different hybrid µEcoG configurations in a modular fashion. To maximize modularity, we designed two linear electrode layouts for this study (10 µm thickness), one with 12 contacts spaced by 0.2 mm and the other with 24 contacts spaced by 0.15 mm, and both with circular, 35 μm diameter platinum recording sites (Figure 2c). Standard cleanroom fabrication processes were used to structure conductive elements on polyimide thin-films using photolithography, resulting in smaller electrodes, traces and higher electrode densities than can be fabricated directly on silicone elastomers^[48,68]^. To achieve a regular 2-dimensional arrangement of recording sites over our cortical sites of interest, polyimide linear arrays were arranged in a regular planar layout on a glass slide and then attached to the underside of a previously fabricated PDMS sheet (Figure 2d). The sheet (0.5-0.8 mm thickness) was placed on top of the polyimide arrays, and PDMS was then applied to the front and the back of the arrays (the locations where no electrode contacts were exposed) to attach them securely to the sheet. Lastly, the metal wire holders were lowered onto the assembly with a micromanipulator, and more PDMS was applied to attach the holders to the PDMS sheet. The transparency of the assembled hybrid µECoG permitted visualization of the underlying brain and of the epifluorescence from populations of cells expressing fluorescent reporters (Fig 2e). We constructed three distinct hybrid µECoGs with different sizes and spatial densities of electrodes.

The first array had 0.15 mm electrode pitch and 0.55 mm strip pitch. It consisted of five linear array strips, each with 24 channels, resulting in a total of 120 electrode contacts and covering an area of 2.2 × 3.45 mm. The second array consisted of ten linear array strips with a pitch of 0.6 mm, each with 24 channels, resulting in a total of 240 electrode contacts and covering an area of 5.4 × 3.45 mm. The third array consisted of 20 linear array strips with a pitch of 0.6 mm, each with 12 channels, resulting in a total of 240 electrode contacts and covering an area of 11.4 × 2.2 mm. The hybrid µECoG designs permitted dense recording from visual cortical areas of the rat, cat and marmoset.

### Hybrid µECoGs record high-quality signals and permit functional mapping from the cortical surface

We first tested the hybrid µECoG in the visual system of the anesthetized cat and rat (Figure 3, cat and Figure S1, rat). The arrays were placed directly on the cortical surface after removal of the dura mater (Figure 3a, Figure S1b) and were able to reliably measure electrical activity from the cortical surface for multiple hours or days within individual terminal experiments under continuous general anesthesia. Arrays were removed and cleaned using detergent and reused in different experiments.

**Figure 3:**
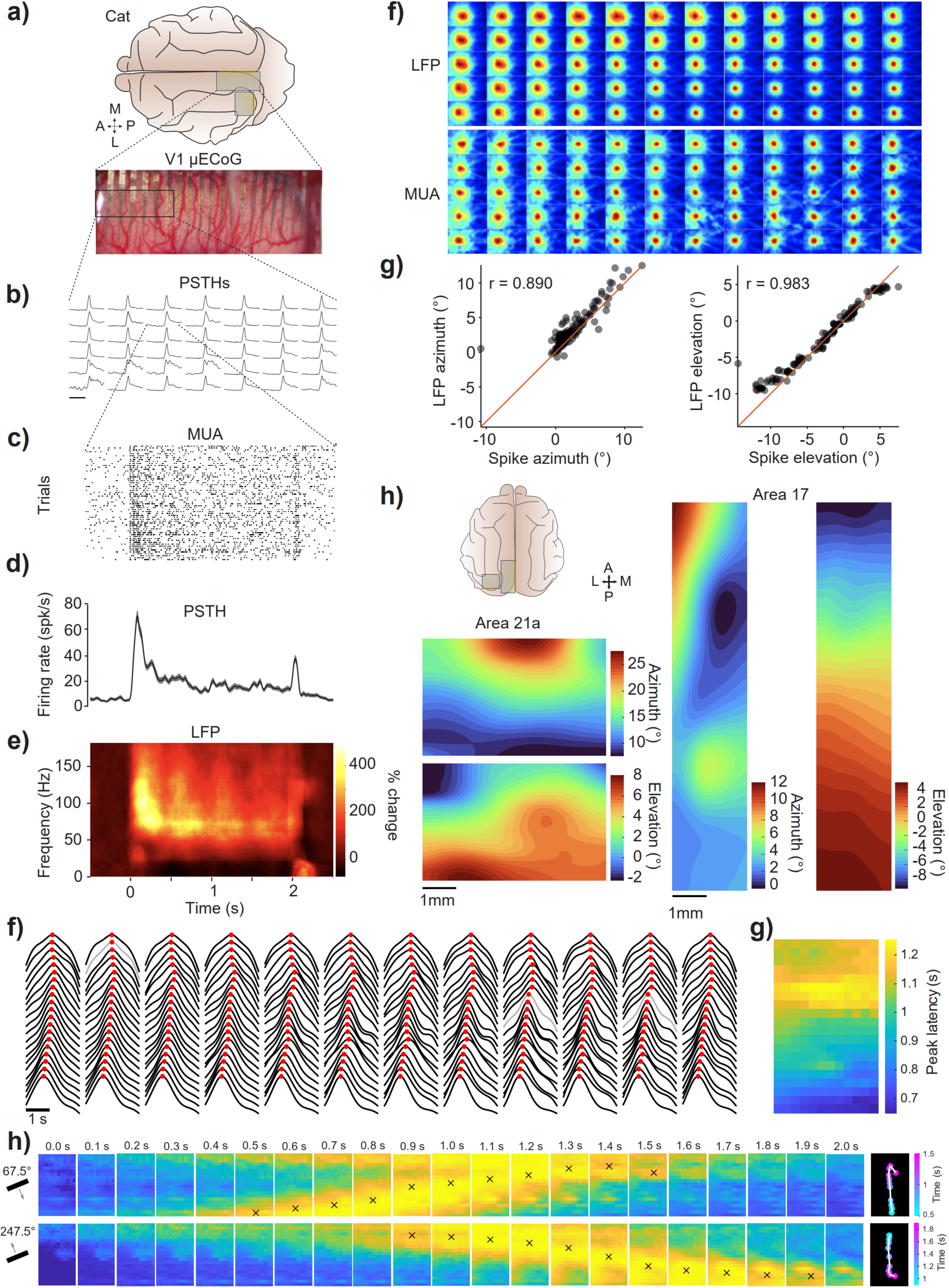
Hybrid µECoG enables dense, multi-scale functional mapping and reveals structured spatiotemporal dynamics of cortical activity in the anesthetized cat. **a)** Illustration of the experimental setup with two µECoG arrays over the cat visual cortex, and photograph of the array over cortical area 17. **a)** PSTHs of the average MUA for a subset of electrodes as indicated by the rectangle in the photograph above. Scale bar equals 500 ms. **c)** Raster plot showing individual MUA spikes detected on one example electrode for multiple trials of visual stimulation. **d)** PSTH and **(e)** time-frequency plot for the full stimulus duration of two seconds. Shaded area in (d) indicates SEM. The x-axis in (e) applies to (c) through (e). The visual stimulus induced strong MUA and sustained gamma-band activity. **f)** Comparison of RFs mapped using gamma-band LFP power or MUA for the same electrode sites in areas 17. **g)** Correlation between the gamma-band LFP and MUA estimated RF center for the azimuth (left) and elevation (right). **h)** Retinotopic maps for cat areas 21a and 17 computed from the LFP-estimated RFs. Retinotopic position varied across cortical areas, consistent with prior electrophysiological and imaging studies. **i)** Average Hilbert envelopes of gamma-band (60–90 Hz) activity for a full-screen moving bar stimulus (67.5° direction), shown for all electrodes in area 17. Red dots indicate detected peak times for each channel. Three noisy channels were interpolated (shown in grey). **j)** Spatial map of response latencies derived from (i), showing a systematic gradient across the cortical surface. **k)** Time-resolved activity maps at successive time points relative to stimulus onset. Upper panels show responses from (i) (67.5° bar stimulus), lower panels show responses for the opposite direction (247.5°). Crosses indicate the location of the detected spatial peak on the array. Rightmost panels show the trajectory of the tracked activation peak over time. For visualization purposes, aspect ratios in panels f–h are not to scale.

To demonstrate the utility of the hybrid µECoG for functional characterization of brain areas, we placed two hybrid µECoG arrays over two anatomically connected and retinotopically organized areas of the cat visual hierarchy. The visual system of the cat is arguably the prototypical system for investigating hierarchical cortical processing and laminar organization.^[69–71]^ To perform large-scale functional characterization, we placed hybrid µECoG arrays over the primary visual cortex (V1, areas 17 and 18) and an intermediate area of the ventral visual stream (area 21a, a homologue of area V4 in the primate^[72]^). We placed one µECoG (20 x 12 electrodes with 0.2 by 0.6 mm spacing) over V1 and a second µECoG (10 x 24 electrodes with 0.15 by 0.6 mm spacing) over area 21a. We found that sites across the µECoG were visually responsive and could record both the aggregate spiking of multiple neurons (multi-unit activity (MUA), Figure 3b-d) and local field potential (LFP) or electroencephalogram (EEG) signals (Figure 3e) from the cortical surface in response to a full-field, full-contrast square-wave grating stimulus. We next performed visual receptive field mapping using a stimulus that consisted of a moving bar (white on black background) moving across the visual field in a random sequence of 16 different directions. Receptive fields could be estimated using either the MUA or the gamma-band LFP activity (Figure S1 for rat and S2 for cat) and resulted in highly consistent selectivity for recording sites (Figure 3f-g). The retinotopic position calculated from the receptive field centers produced functional maps that revealed the expected retinotopic organization of visual areas in the cat (Figure 3h). The visual receptive fields in area 21a appeared on average larger and moved more in visual space in a smaller amount of cortical space than those in area 17.

We next inserted intracortical microelectrodes through the PDMS portion of the hybrid µECoG, adjacent to the polyimide thin-film linear electrode arrays to perform simultaneous depth recordings from functionally characterized neuronal populations, first with tungsten microelectrodes and subsequently with intracortical multi-electrode arrays. The retinotopic selectivity of neurons recorded with penetrating tungsten electrodes from the underlying cortex matched well with that expected based on the functional mapping with the hybrid µECoG (Figure S2b-c, right). This close correspondence enabled targeted recordings of functionally characterized populations within an area of interest. Moreover, the placement of multiple µECoG arrays permitted the simultaneous mapping of visual space in area 17 and area 21a and the subsequent targeting of retinotopically overlapping neurons across the cortical depth simultaneously in both areas (Figure S2b-c). The functional map estimated with the µECoG allowed us to target portions of area 17 and 21a representing similar portions of visual space and likely to be anatomically connected.

### Hybrid µECoGs enable multi-scale investigation of cortical activity

To demonstrate the capacity for multi-scale investigation of cortical activity patterns both topographically across the cortical surface, but also in the depth with laminar electrode arrays, we combined the surface recordings from the µEcoG in cat area 21a with a multi-shank laminar electrode array (Figure 4a). We inserted a 128-channel electrode array (Atlas Neurotech, 4 shanks with each 32 electrodes spaced by 100 µm) based on functional mapping at the desired retinotopic position at the medial edge of the µEcoG array (Figure 4b). The combined recording enables correlating the 2D surface signals from the µEcoG with the signals from the laminar array across all depths of the cortex. The topography of the resulting correlation values depended on the frequency band of the cortical activity, the depth along the laminar electrode array, as well as the position of the laminar electrode array relative to the µEcoG (Figure 4c). The correlation pattern from the high frequency activity (100-200 Hz) showed a singular localized spatial peak at the position closest to the corresponding laminar array (colored arrows in Figure 4c). Specifically, the correlation peak for shank 1 was located at the medial edge of the array, slightly posterior from the center, whereas the peak for shank 4 was located at the most posterior medial edge, as expected from the location of the laminar arrays (Figure 4b). Interestingly, the sign of the localized correlation peaks flipped from positive to negative deeper in the brain. This reversal in correlation sign is consistent with the known reversal of the LFP across layers.^[73]^ The correlation patterns from the low frequency LFP components (10-20Hz) resembled this pattern but it was less localized and showed larger absolute correlation values, as well as additional spatial features when compared to the high frequency patterns. These frequency-dependent differences likely reflect the spatial scale of the underlying neural signals. Low-frequency LFP components primarily reflect synaptic inputs integrated over larger neuronal populations and therefore spread over broader cortical distances and exhibit millimeter-scale spatial correlations, whereas higher-frequency activity is more closely related to local population spiking and synaptic currents and thus has a more spatially restricted footprint.^[74]^

**Figure 4:**
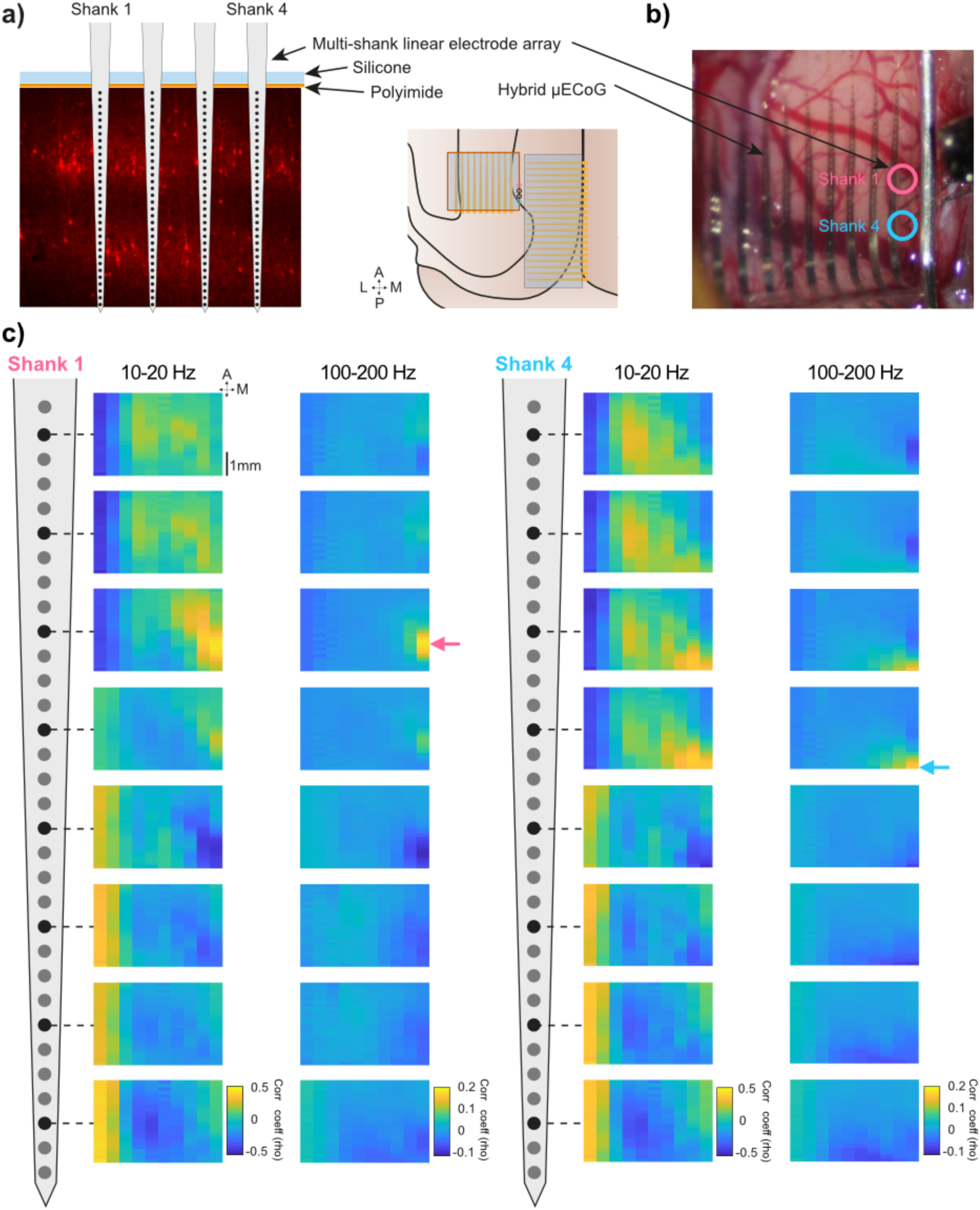
Hybrid µECoG enables investigation of multi-scale interactions between laminar and topographical activity patterns. **a)** Schematic illustration of a multi-shank laminar electrode array inserted at the edge of a hybrid µECoG in area 21a based on retinotopic functional mapping. **b)** Photograph of the high-density hybrid µECoG placed over area 21a with the inserted multi-shank laminar electrode array. Circles indicate positions of shanks 1 and 4. **c)** Topographical pattern of correlation between two different shanks of the laminar array and the band-limited LFP in two frequency ranges showing differential 2D correlation patterns by frequency and depth. Note the singular localized spatial peak at the position closest to the corresponding laminar array.

### Hybrid µECoG for functionally targeted multi-area laminar recordings in the marmoset

To aid the investigation of inter-areal interactions in the visual system, we next used the hybrid µECoG to map multiple areas in the visual hierarchy of the anesthetized marmoset. We placed two µECoG arrays over the primary visual cortex (V1) and an intermediate visual area (medial temporal area, MT/V5) to map their retinotopic organization. We then used the µECoG-derived retinotopic maps to guide the insertion of linear depth electrodes for functionally aligned multi-area laminar recordings (Figure 5a). The placement of the laminar probe in MT was guided by the retinotopic map to approximate RF alignment with the V1 site. Additionally, visible blood vessels were avoided during insertion. The soft PDMS encapsulation allowed repeated penetrations with laminar probes, resealing after each penetration and retraction (Figure S7). As with the rat and cat, we were able to estimate visual receptive fields from the LFP and MUA recorded from the surface of the brain in the marmoset (Figure 5b). Receptive fields recorded from the laminar multi-electrode arrays inserted through the hybrid µECoG matched those of nearby electrodes on the µECoG array but not those of more distant electrodes (Figure 5b). This allowed simultaneous laminar recordings of targeted regions in areas V1 and MT of the marmoset. The median RF distance between the V1 and MT laminar probes was 1.54° (95% CI, 1.11–23.98), consistent with close retinotopic alignment resulting from µECoG-guided targeted insertion of the MT probe. To estimate the RF distance expected if the MT probe had been inserted randomly within the MT array, we performed a Monte Carlo simulation based on the empirical RF location of the V1 probe and the retinotopic map derived from the RF centers of the MT µECoG electrodes (Figure S4). Random MT penetrations yielded an expected RF distance of 4.97 ± 3.75° (mean ± SD), compared to the observed V1–MT RF distance of 1.54°. Thus, the actual penetration was better aligned than 91.3% of random penetrations. These results demonstrate that µECoG-guided targeting substantially improves the alignment of RF locations across cortical areas.

**Figure 5:**
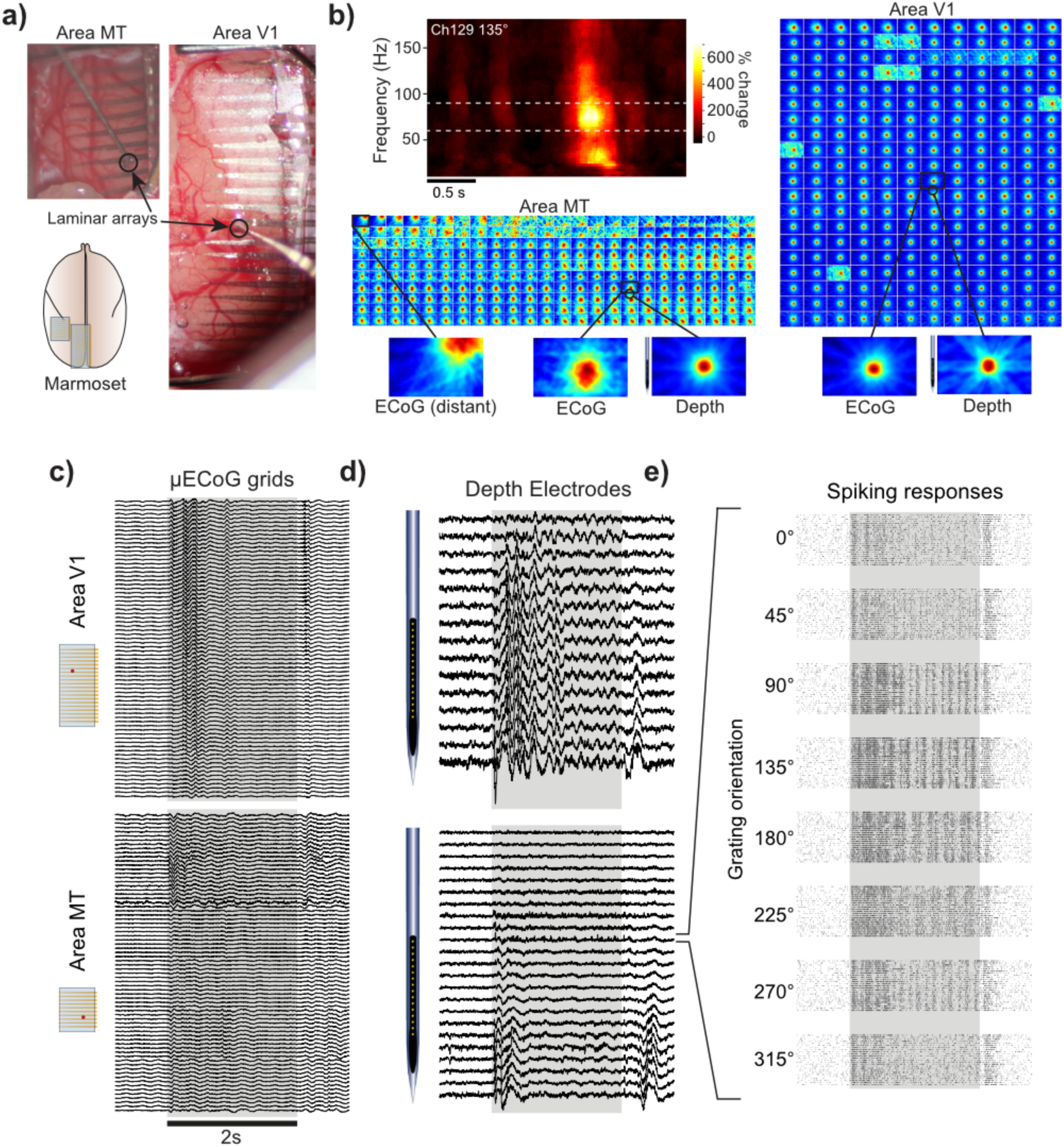
Hybrid µECoG enables functionally targeted multi-area laminar recordings in the anesthetized marmoset. **a)** Two hybrid µECoG arrays placed over area V1 and MT in the marmoset visual cortex to guide the insertion of laminar arrays for paired multi-area laminar recordings in the primate visual hierarchy. **b)** µECoG electrodes demonstrated visual responsiveness and visual mapping could be performed to determine retinotopic organization across cortical areas. The retinotopic selectivity of neurons recorded from the depth matched well the selectivity expected based on surface recordings with the µEcoG as evident by the similarity of receptive fields from the depth probe and the neighboring µEcoG sites. **c)** Example single-trial LFP responses from a subset of simultaneously recorded µECoG channels. In the example trial shown, a full screen drifting grating was presented for a duration of 2 s. The onset and offset of the visually evoked potentials are visible in the traces from area V1 and MT and **d)** across the depth of the cortex simultaneously recorded from the laminar arrays in both areas. **e)** Example raster plots of spiking activity recorded from the cortical depth with a laminar array in MT. Each block shows multiple trials of visual stimulation with a particular stimulus orientation, demonstrating the ability of detailed functional characterization of the neuronal populations underlying the µECoG.

In V1, receptive field centers appeared relatively similar across neighboring µECoG channels. This reflects the strong cortical magnification of central visual space in primate V1, where large regions of cortex represent small portions of visual space, such that electrodes separated by hundreds of micrometers can exhibit similar receptive field locations. In contrast, receptive field locations changed more rapidly across the cortical surface in MT, consistent with the more compressed retinotopic organization of higher visual areas. Consequently, µECoG-based retinotopic mapping provides less spatial specificity in V1 than in higher visual areas. At the same time, this highlights the challenge in areas such as MT of predicting, based on anatomy alone, where to insert electrodes to obtain overlapping receptive fields with V1, and underscores the utility of our approach. Simultaneous laminar and µECoG recordings permit activity across the cortical depth to be related to ongoing activity in local and distant cortical populations according to their functional correspondence (Figure 5c-d). Spiking activity recorded from the cortical depth also demonstrated the known orientation and direction selectivity of cortical columns in the primate and cat primary visual cortex (Figure 5e). Such an approach can greatly increase the efficiency of localizing functionally aligned populations for multi-area laminar recordings. Furthermore, simultaneous µECoG and laminar recordings allow the investigation of cortical interactions across spatial scales.

### Hybrid µECoG for optogenetic mapping of local and inter-areal interactions

We next took advantage of the optical access through the hybrid µECoG to perform local and inter-areal optogenetic mapping. Prior to the recordings, cats and marmosets were injected with adeno-associated viral (AAV) vectors to express channelrhodopsin in cortical pyramidal neurons (CamKII promoter). After multiple weeks of expression, experiments were performed, in which we positioned an optic fiber connected to a laser over the surface of the µECoG to permit direct stimulation of neurons lying below. As previously mentioned, the µECoG permitted the localization of populations expressing the transgene due to the attached fluorophore (Figure 2e). Visualization and widefield fluorescence microscopy of the cortical surface through the µECoG enabled us to directly target cortical populations expressing the excitatory opsin while avoiding large vasculature (Figure 2e, Figure 5a and Figure 6a).

**Figure 6:**
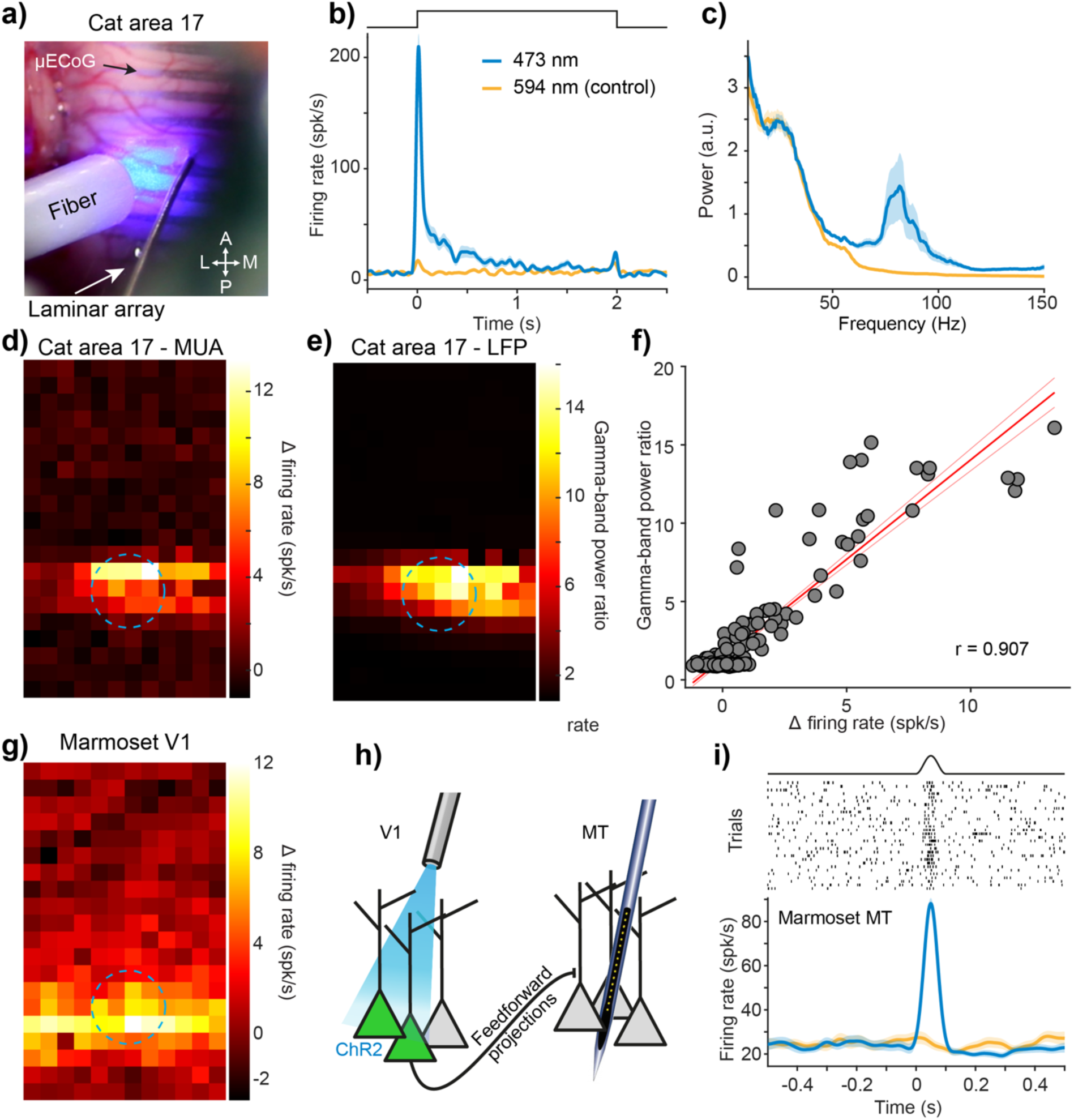
Hybrid µECoG permits local and interareal optogenetic mapping in the visual hierarchy. **a)** Photograph of a hybrid µECoG placed over area 17 of the anesthetized cat during simultaneous multi-area laminar recording and optogenetic stimulation of pyramidal neurons. **b)** A peristimulus time histogram of MUA activity recorded on the µECoG during optogenetic activation of neurons in area 17 (blue) or control (yellow). **c)** Power spectrum of LFP power recorded on the µECoG probe in area 17 to local optogenetic activation (blue) or control (yellow). **d)** Spatial response pattern of MUA activation over area 17 to focal optogenetic activation. **e)** Spatial response pattern of gamma-band power increase over area 17 to focal optogenetic activation. Dashed blue circles in (d) and (e) indicate estimated laser location. **f)** Local increases in MUA and gamma-band power correlate across the µECoG in area V1. **g)** Spatial response pattern of MUA activation over marmoset V1 to focal optogenetic activation. **h)** Illustration of optogenetic feed-forward neuronal activation of neurons projecting from area V1 to area MT as performed in our study. **i)** Example channel MUA activity evoked in area MT to feedforward optogenetic activation of V1 neurons. MT laminar probe was inserted based on functional mapping of area MT to retinotopically correspond to the V1 optogenetic stimulation site.

In cat area 17, focal optogenetic activation of a local population of excitatory neurons expressing an excitatory opsin (hChR2 under the CamKII promoter) increased neuronal activity detectable in both MUA and gamma-band LFP power on the µEcoG array (Figure 6b and c). The topographical pattern of this local activation could also be visualized across the cortical surface (Figure 6d and e). Optogenetically activated neuronal activity in area 17 was localized to a focal area consistent with the location of the laser, and increased MUA correlated with increased gamma-band power as detected on the µECoG (Figure 6f). Similarly, in marmoset V1, optogenetic activation also led to spatially localized MUA increase around the location of the laser (Figure 6g). Feedforward projections from V1 to MT have been well established in non-human primates, including marmosets.^[75–78]^ Thus, activation of neurons in V1 is expected to evoke responses in MT, either directly (Fig. 6h) or via intermediate visual areas such as V2. Indeed, we observed that stimulation with blue light, but not yellow control light, caused short-latency responses in MT (Fig. 6i; Fig. S5b; 8 out of 32 channels showed significant modulation; two-sided signrank test, Bonferroni corrected; p-range = 3.71e-30–1.20e-05). Since feedforward projections are expected to be specific to matching retinotopic locations in the downstream area, we analyzed whether the

optogenetic responses depended on the retinotopic alignment of the MT laminar probe with the stimulated V1 location. The retinotopic location of optogenetic stimulation was defined by the median RF location of the V1 laminar probe at the stimulation site. In the recordings shown in Fig. 6i, a second laminar probe was inserted through the soft array into MT. The median RF location of the first MT probe was closer to the V1 probe (1.56°, 95% CI, 1.13–3.66°), whereas the second probe was further away (4.08°, 95% CI, 2.25–6.30°; permutation test, p = 0.012; Fig. S5 b). Optogenetically evoked feedforward spiking responses were detected only on the MT probe with the smaller RF distance to the V1 probe, and thus closer to the retinotopic location of the optogenetic stimulation (Fig. S5c). No responses were observed on the more distant MT probe (Fig. S5d).

Notably, we did not find evidence for optogenetically induced feedforward activity in the MUA of the µECoG in MT (Figure S5e, no channels showed significant modulation; two-sided signrank test, Bonferroni corrected). This may reflect the limited sensitivity of surface µECoG recordings, which primarily capture population activity from superficial cortical layers. In contrast, the relatively weak feedforward activation from V1 may predominantly reach the input layers of MT and therefore remain below the detection threshold of the surface electrodes.

### µECoG enables large-scale recording across the cat visual hierarchy

Finally, we sought to perform large-scale recordings of activity across the visual hierarchy of the cat using a combination of our hybrid µECoG and a commercially available µECoG developed with an industry partner (CorTec GmbH, Freiburg, Germany). To this end, we worked with CorTec GmbH to design and fabricate a lower-density pure PDMS µECoG array with 120 electrodes (0.2 mm diameter platinum-iridium electrode sites) arranged in a regular 6 x 20 electrode grid with 0.7 mm by 0.8 mm pitch (Figure S3a and b). The overall dimensions of the CorTec array were 4.5 mm by 16.2 mm, and it could be placed to cover multiple visual areas in the cat. Like our µECoG, the CorTec µECoG was transparent (Figure 7a and b) and could be placed to cover multiple areas in the cat visual cortex (Figure 7c). We placed the CorTec µECoG to cover the primary visual cortex of the cat (areas 17 and 18) and performed functional mapping (retinotopy) of the underlying cortex. LFP signals recorded from the CorTec µECoG could be used to determine the visual selectivity of the underlying cortex, nicely demonstrating the difference in receptive field size when moving from area 17 to area 18 (Figure 7d). Furthermore, as with our hybrid µECoG, linear electrode arrays could be inserted to perform simultaneous functionally guided laminar recordings (Figure 7b). We therefore attempted to scale up our µECoG-based recording of activity across the visual hierarchy of the cat. Using 2 hybrid µECoG arrays and 5 silicone µECoG arrays, we were able to cover large portions of the dorsal cortex of the cat bilaterally. We were able to record simultaneous activity from 7 µECoG arrays totaling 1080 electrodes across 9 visual areas bilaterally (Figure 7e,f). Example traces from the simultaneous µECoG recordings highlight the possibility to investigate high-resolution spatiotemporal dynamics across extended portions of the cortical hierarchy (Figure 7f,g).

**Figure 7:**
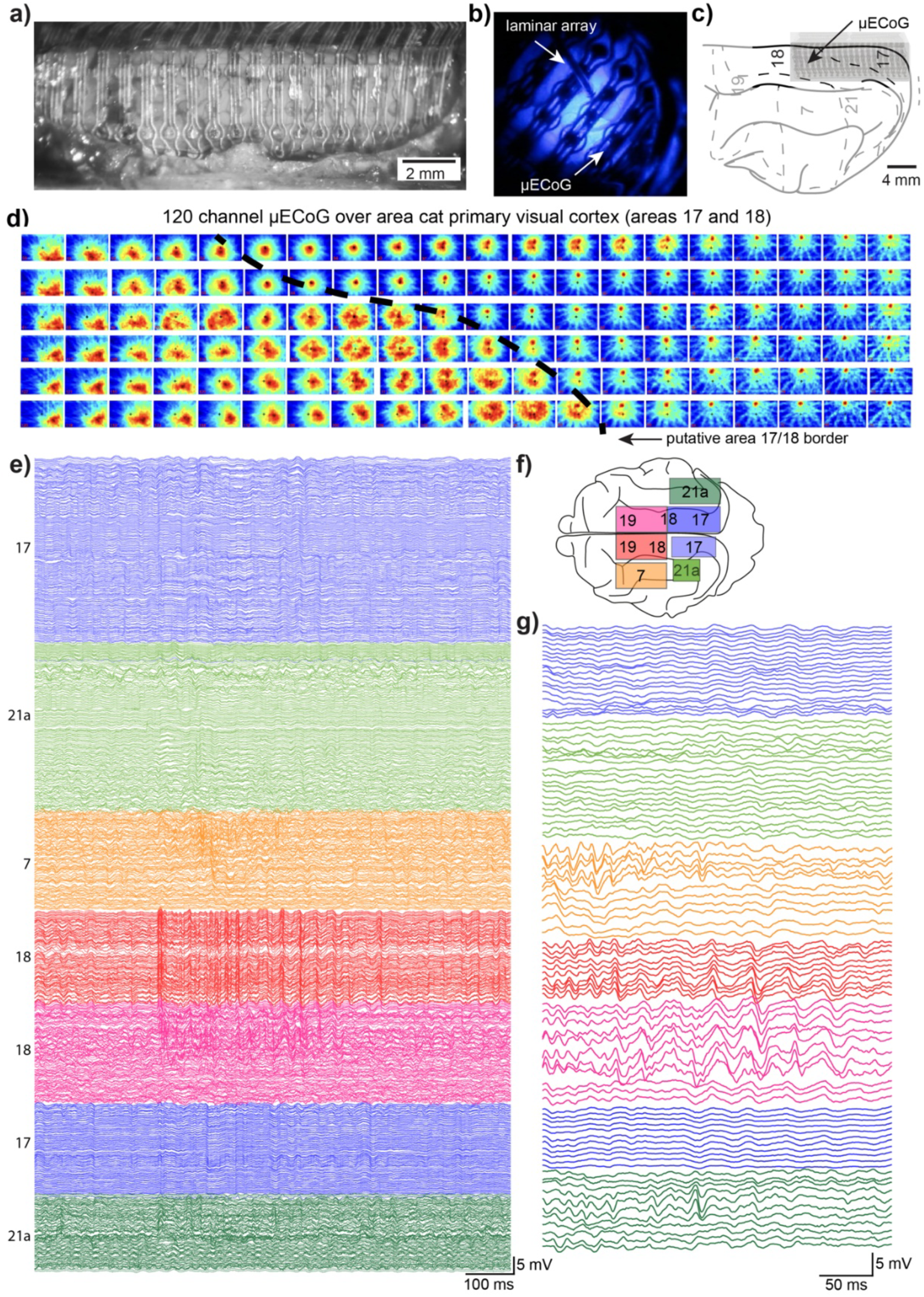
µECoG enables dense, multi-scale functional mapping of cortical activity across the visual system in the anesthetized cat. **a)** Photograph of a CorTeC µECoG in place over the cat primary visual cortex (areas 17 and 18). **b)** Photograph of blue excitation light from a laminar electrode array with integrated fiber-optics, transmitting through the cat visual cortex. **c)** Illustration of the µECoG array as targeted to the cat cortical atlas using anatomical coordinates to target primary visual cortex. **d)** Visual receptive fields recorded with the CorTec µECoG with the putative border between area 17 and 18 illustrated and visible in the larger sized receptive fields in area 18. **e)** 1080 channels of data recorded using two custom hybrid µECoG arrays and five commercial µECoG arrays across the visual system of the cat. **f)** Illustration of the placement of individual µECoG arrays over the dorsal cortex of the cat. **g)** Zoom-in on a selection of sites from each µECoG array showing fluctuations in the LFP recorded across visual areas.

## Discussion

Brain activity displays exquisite coordination across spatial and temporal scales. New tools are required to simultaneously observe and manipulate multiscale brain activity, especially from anatomically connected populations distributed across brain-wide networks. We present a hybrid µECoG that enables functionally targeted multiscale investigations of local and distributed neural populations. We combined the advantages of two key, biocompatible flexible polymers to produce an ECoG array that had high electrode density, while conforming to soft neural tissue. By combining flexible, thin-film electrode arrays based on polyimide with a silicone elastomer substrate, we constructed a series of modular µECoG arrays that provided high-resolution recordings from distributed cortical brain areas. The two-step fabrication process isolated demanding clean-room fabrication from flexible, modular assembly of the final arrays to match different cortical areas of interest across the widely varying brain sizes of rats, cat and marmosets. We demonstrate functionally guided multi-area laminar recordings and inter-areal optogenetic activation to highlight the utility of this approach for investigating communication and coordination between distributed neuronal populations.

Our application of the hybrid µECoG demonstrates their utility for investigating the multi-scale and multi-area characterization and for probing causal inter-areal responses using targeted optogenetic stimulation. Despite reaching these goals, all our experiments were performed in anesthetized animals without the need to package the arrays in a robust fashion conducive to long-term chronic recording in awake, behaving animals. Within the experiments presented here, arrays could be reused across multiple acute experiments that each lasted up to several days, without evidence of strong degradation in noise levels and only minor changes in impedance (Supplemental Notes, Fig. S6). However, this assessment is limited to repeated acute use and does not capture long-term stability under chronic implantation conditions. Thus, systematic evaluation of long-term recording stability in chronic preparations should be addressed in future work. Nevertheless, similar arrays have already been integrated with recording chambers and artificial dura replacement for chronic implantation in primates or cats^[79–82]^, and related designs have been chronically implanted in mice for months.^[51,83]^ Indeed, our use of silicone elastomers was inspired by their previous use as chronic dura-replacements to achieve long-term optical and electrical access to the cortical surface.^[59,66,67]^ Here we extend that use to enable functional mapping across multiple cortical areas, while permitting repeated targeted recording with penetrating electrodes and laminar arrays. Our design enabled simultaneous recording from neuronal populations in the depth due to the soft PDMS encapsulation that could be penetrated with stiff, linear electrode arrays, resealing after penetration. The optical transparency of PDMS provided visual access to the cortical surface. This enabled identification of regions with strong opsin expression, optogenetic stimulation with a surface light source and helped avoid blood vessels during electrode placement. In future studies, this transparency might also support optical recordings to investigate cellular and subcellular mechanisms underlying local and interareal interactions.^[16,51]^

While optical imaging was not evaluated in the present study, inter-areal investigations could, in principle, be conducted in an all-optical manner by co-expressing genetically encoded activity indicators and excitatory or inhibitory opsins in defined neuronal populations. Such methods have been impressively demonstrated in the mouse^[84–87]^, they have not yet been achieved in the larger brains of the cat or primate. Although the simultaneous use of 2-photon imaging and optogenetic stimulation has recently been achieved in primates within a single cortical area,^[88–92]^ the transfer of appropriate viruses and molecular tools, as well as suitable optical systems to study multiple areas in the considerably larger feline or primate brain is non-trivial. Considerable difficulties may arise due to challenges with viral co-infection, larger distances between relevant areas and increased brain movement in larger animals. That being said, optical methods can enable higher specificity by enabling viral and genetic approaches to target specific populations and project pathways. However, even the most advanced optical methods, using genetically encoded voltage indicators cannot detect single spikes with millisecond precision across populations that span several millimeters.^[93]^ Therefore, we believe that the combined use of optical and electrical recording methods provides the opportunity for synergistic, multimodal investigations of brain function.^[16,51]^ Future work could establish such hybrid µECoG arrays as a platform for chronic brain interfaces to enable long-term multi-scale optical and electrophysiological recording of brain activity.

While functional mapping with µECoG and subsequent local recording with multi-electrode arrays can be performed sequentially, the combined application of large-scale and dense measurement methodologies simultaneously permits local cellular activity to be evaluated in the context of ongoing patterns of brain wide activity with which it is coordinated.^[15,16,25]^ Such investigations have begun to reveal the importance of multi-scale studies for interpreting the responses of single cells in the context of local circuit activity and situating local patterns in global brain states.^[9,25,31,32,94,95]^ Functional mapping also permits targeted injection of viruses or pharmacological agents, further expanding the possibilities of combined µECoG and depth electrode or optical studies. For example, viral transfection of opto- or chemogenetic effectors into local or long-range projections may permit functionally characterized populations to be activated or silenced, expanding the possibilities to dissect inter-areal interactions^[89,90][93,94][94,95][96,97]^, including feedback effects from higher areas (Nurminen et al. 2018, Lewis et al. 2024). Likewise, viral transfection of genetically encoded activity indicators can permit simultaneous optical recording of cellular signals in the local population or identified projections, such as long-range neuronal or neuromodulatory projections that might affect local or inter-areal dynamics.^[31,98–101]^ The combination of electrical recording with optical methods, such as imaging,^[16,94,102]^ or optical perturbations^[27,96,103,104]^ can permit fined-grained investigations that further reveal the cellular processes underlying coordinated dynamics and inter-areal interactions. New techniques and tools to measure and manipulate brain activity have revolutionized our understanding of healthy and disordered brain function. The hybrid µECoG we developed enables the combination of simultaneous optical and electrophysiological measurement from functionally characterized local and distributed neuronal populations. Such tools and the studies they enable will allow refined investigation of inter-areal information transfer and the local and distributed processes underlying adaptive behavior.

## Methods

### ECoG fabrication

The hybrid µECoG arrays were fabricated in a two-stage process. First, flexible, linear arrays were fabricated using photolithography to structure platinum on polyimide thin films. Second, these arrays were arranged in a regular pattern and bonded to a PDMS superstructure.

#### Fabrication of flexible linear arrays

Using standard MEMS cleanroom techniques as previously published,^[48,63,65]^ we fabricated 12 or 24 contact linear arrays on polyimide with 35 µm diameter platinum contacts. Briefly, we spin-coated a 5 μm layer of polyimide (U-Varnish S, UBE) on a silicon wafer and imidized it in a vacuum oven at 450 °C. Next, activation was performed with O2-plasma (30 s, 100 W), and we sputtered a 300 nm layer of platinum onto the polyimide and defined electrodes and tracks with photolithography. The metal tracks were insulated with a second layer of polyimide (5 μm). Finally, the electrode pads and the overall linear array shape were defined via reactive ion etching in an O2-plasma.

#### Integration with PDMS

The previously fabricated linear arrays were arranged into the desired pattern and then integrated to the PDMS superstructure. First, we arranged the linear arrays into a regular 2D sheet by placing them with the exposed electrode surface facing down onto a clean glass slide that was placed over millimeter grid-paper. Under stereoscope guidance and with the aid of the grid-paper, we aligned the polyimide linear arrays by hand to the desired spacing and coverage. To prevent accidental misalignment of the arrays or unintended occlusion of electrode contacts, we did not pour fluid PDMS on top of the polyimide strips for in-situ polymerization. Rather, a previously polymerized PDMS sheet (0.5-0.8mm thickness) was cut to the desired size and then placed on top of the polyimide arrays. Next, any misalignment was manually corrected, and PDMS was applied to the front and the back of the arrays. This prevented any leakage of liquid PDMS onto the electrode contacts, which could lead to impedance increase or full electrical insulation, thereby rendering them non-functional. Lastly, the metal wire holders were lowered onto the assembly with a manual micromanipulator and more PDMS was applied to attach the holders to the PDMS sheet. We used a two-component PDMS (Biopor AB, 25 Shore) for all manufacturing steps. PDMS curing was performed at room temperature.

### Animals

Two adult male Sprague-Dawley rats (Rattus norvegicus), three adult domestic cats (*Felis catus*; 2 females), and two adult male marmosets (Callithrix jacchus) were used in this study. We used multiple organisms to establish the utility of the approach across diverse model species. All procedures complied with the German law for the protection of animals and were approved by the regional authority (Regierungspräsidium Darmstadt). After an initial surgery for the injection of viral vectors and a 4-6 week period for opsin expression, all recordings were obtained during terminal experiments under general anesthesia without recovery or chronic implantation.

Recordings from cat visual cortex (Figs. 2, 3, 4, 6, 7) spanned approximately 136.6 hours (∼5.7 days) under continuous general anesthesia. Recordings from marmoset visual cortex (Figs. 5, 6) spanned approximately 82.6 hours (∼3.4 days) under continuous general anesthesia. Throughout terminal recording experiments in marmosets and cats, physiological parameters were continuously monitored and actively maintained within normal ranges.^[96,105]^ Body temperature was controlled using a feedback-regulated heating system, and respiration was controlled via artificial ventilation. Anesthetic depth and ventilation parameters were adjusted as needed to maintain stable end-tidal CO₂, heart rate, oxygen saturation, and body temperature.

### Viral vector injection

#### Cats

Anesthesia was induced by intramuscular (i.m.) injection of ketamine (10 mg/kg) and dexmedetomidine (0.02 mg/kg). Cats were intubated and anesthesia was maintained with N2O:O2 (60/40%), isoflurane (∼1.5-2.5%) and remifentanil (0.3 µg/kg/min). Rectangular craniotomies were performed with an ultrasonic bone drill (Piezosurgery, Mectron) over area 17 and the dura mater was removed. The target location in area 17 was guided by stereotaxic coordinates^[106]^ and by the local pattern of sulci and gyri. Three to four injection sites were chosen, avoiding blood vessels, with horizontal distances between injection sites of at least 1 mm. At each site, a Hamilton syringe (34G or 35G needle size; NanoFil syringe, World Precision Instruments) was inserted with the use of a micromanipulator (David Kopf Instruments) and under visual inspection to a cortical depth of 1 mm below the pia mater (0.5mm below pia in the rat). Subsequently, 2 µl of viral vector dispersion was injected at a rate of 150 nl/min with a microinjector pump (UMP3-1, World Precision Instruments). After each injection, the needle was left in place for 10 min before withdrawal, to avoid reflux. Upon completion of injections, the dura opening was covered with silicone foil and a thin layer of silicone gel, the trepanation was filled with dental acrylic, and the scalp was sutured.

In all cats, area 17 of the left hemisphere was injected with AAV1-CamKIIα-hChR2(H134R)-eYFP (titer: 1.22*10^13^ GC/ml).

#### Marmosets

Anesthesia was induced via i.m. injection of alfaxalone (8.75 mg/kg) and diazepam (0.625 mg/kg), then maintained with isoflurane (∼1.5–2.5%) delivered through a nose cone in one animal and through an endotracheal tube in the other animal. Buprenorphine (0.05 mg/kg) was administered pre-surgery for analgesia. A small burr hole (∼3 mm) was drilled over the target site in the left primary visual cortex guided by stereotaxic coordinates^[107]^ and the dura was carefully incised using the bent tip of a hypodermic needle. A Hamilton syringe with a 35G needle (NanoFil syringe, World Precision Instruments), mounted on a micromanipulator (David Kopf Instruments) was then slowly lowered into the brain. One animal received injections at two sites within the same craniotomy at depths of 0.8 mm and 0.5 mm, with 2 μL delivered per site. The other animal was injected at a single site with 1.6 μL. In both animals, AAV1-CamKIIα-hChR2(H134R)-eYFP viral vectors were injected at 200 nL/min. After injection, the trepanation was covered with dental acrylic, and the scalp was sutured.

#### Rats

Anesthesia was induced by i.m. injection of ketamine (100 mg/kg) and dexmedetomidine (0.25 mg/kg), then maintained with isoflurane (∼1.5–2.5%) in 100% oxygen delivered through a nose cone. The viral vector injection was performed in the primary visual cortex^[108]^ and the procedure was otherwise identical to that described for the marmoset.

All injections were performed under aseptic conditions for rats and under sterile conditions for marmosets and cats. The DNA plasmids were provided by Dr. Karl Deisseroth (Stanford University, Stanford, CA). AAV1 and AAV9 viral vectors were obtained from Penn Vector Core (Perelman School of Medicine, University of Pennsylvania, USA).

### Neurophysiological recordings - cat

For recording experiments, anesthesia was induced and initially maintained as during the injection surgery, only replacing intubation with tracheotomy and remifentanyl with sufentanil. After surgery, during recordings, isoflurane concentration was lowered to 0.6%-1.0%, eye lid closure reflex was tested to verify narcosis, and vecuronium (0.25mg/kg/h i.v.) was added for paralysis during recordings to prevent reflexive eye movements. Throughout surgery and recordings, Ringer solution plus 10% glucose was given (20 ml/h during surgery; 7 ml/h during recordings), and vital parameters such as ECG, body temperature and end-tidal CO2 were monitored continuously.

### Neurophysiological recordings - marmoset

For recording experiments, anesthesia was induced and initially maintained as during the injection surgery. A tracheotomy was performed and sufentanil (4-8 µg/kg/h) was continuously infused intravenously in Ringer solution containing 5% amino acids and 3% glucose. After surgery, during recordings, isoflurane concentration was lowered to 0.5%-1.5%, eye lid closure reflex was tested to verify narcosis, and vecuronium (0.2 mg/kg/h) was added for paralysis during recordings to prevent reflexive eye movements. Throughout surgery and recordings, vital parameters such as ECG, body temperature and end-tidal CO2 were monitored continuously.

### Neurophysiological recordings - rat

For recording experiments, anesthesia was induced and maintained as during the injection surgery.

### Neurophysiological recordings - general

Each experiment consisted of multiple recording blocks of data acquisition per animal. Large rectangular craniotomies were made and the dura was removed to permit placement of µECoG arrays directly on the brain surface in the respective areas. For depth recordings, several electrode types were inserted though the µECoG: Tungsten microelectrodes (∼1 MΩ impedance at 1 kHz; FHC), 32- or 128-contact silicon probes (50 or 100 µm inter-contact spacing; NeuroNexus or ATLAS Neuroengineering), and 16- or 24-contact linear electrode arrays (V- and U-probes; Plexon). In one cat, one 16-contact probe with 150 µm inter-contact spacing and one 46 µm optic fiber, and one 16-contact probe with 150 µm inter-contact spacing and four 46 µm optic fibers were used (Plexon V- and U-probe, respectively). Neural signals were recorded with the OpenEx suite software package (Tucker Davis Technologies) through active, unity gain head stages (ZC32, ZC64 and ZC128, Tucker Davis Technologies) and digitized at 24,414.0625 Hz (PZ2 preamplifier, Tucker Davis Technologies). For MUA recordings, the signals were filtered with a passband of 700 to 7000 Hz, and thresholds were set to retain the spike times of small clusters of units. For LFP recordings, the signals were filtered with a passband of 0.7 to 250 Hz, digitized at 1017.1 Hz and resampled to 1 kHz for analysis. After each experiment, µECoG arrays were cleaned by soaking them overnight in enzymatic detergent (Tergazyme) and then rinsed with DI water and stored dry.

### Visual stimulation

Visual stimuli were displayed on a gamma-corrected TFT monitor (Samsung SyncMaster 2233RZ) at a refresh rate of 120 Hz and generated via custom MATLAB (MathWorks) code and Psychtoolbox-3.^[109]^ In cats and marmosets, contact lenses were used to prevent drying of the cornea and to assure correct focus of the visual stimuli onto the retina.

### Optogenetic stimulation

Optogenetic stimulation was performed with a 473 nm (blue) laser or with a 470 nm (blue) LED (Omicron Laserage). A 594 nm (yellow) laser was used as randomly interleaved control. Laser light was delivered to the cortical surface through a 100 µm or a 200 µm diameter multimode fiber (Thorlabs), LED light through a 2 mm diameter polymer optical fiber (Omicron Laserage). Fiber endings were placed just above the cortical surface, immediately next to the recording sites with a slight angle relative to the electrodes. Laser waveform generation used custom circuits in TDT, and timing control used Psychtoolbox-3, a toolbox in MATLAB.

### Data analysis

All data analysis was performed using custom code and the Fieldtrip toolbox,^[110]^ both written in MATLAB (MathWorks). PSTH plots for MUA rates were smoothed with a Gaussian window (SD = 10 ms). For receptive field mapping analysis from LFPs, signals were band-pass filtered with a Butterworth filter between 60 and 90Hz (filter order = 2). The absolute amplitude of the Hilbert transform of this signal was then used to generate a trial averaged, smoothed PSTH (SD = 60 ms) for each of the 16 directions of the moving bar. The value at a delay of 50 ms was used as the magnitude of visual response at the corresponding location in visual space and the 2D receptive field image was spatially smoothed (SD = 25.5 pixels). Receptive field mapping analysis from MUA data was identical, except that the PSTH was calculated from smoothed spiking data (SD = 60 ms).

The center of each receptive field was calculated by thresholding the normalized 2D receptive field image data at a value of 0.7 and then calculating the center of mass. Retinotopic maps for the full areas covered by the µECoGs where then generated from the resulting receptive field centers. The map was upscaled by a factor of 50 using bicubic interpolation and smoothed with a Gaussian filter (SD = 50 pixels). For the maps from cat area 21a (Figure 3h), 8 out of 240 channels were excluded and interpolated because they appeared electrically shortened, as indicated by an abnormally high correlation with other channels (defined as a Pearson correlation >0.97 with at least 8 other channels). For the correlation of LFP-based with spike-based receptive field coordinates, we only included channels with a signal-to-noise ratio (SNR) > 2.5. SNR was defined as the ratio of the pixel values inside the normalized receptive field (all pixels that were >0.7) and the rest of the image, resulting in 138 out of 240 channels being included in the analysis (Figure 3g).

LFP time-frequency analysis was calculated for data epochs that were adjusted for each frequency to have a length of 4 cycles and moved over the data in a sliding-window fashion in 1-ms steps. Each epoch was multiplied with a Hann taper, Fourier transformed, squared and divided by the window length to obtain power density per frequency. For the different stimulation frequencies f, LFP power is shown as percent change of power during stimulation versus pre-stimulation baseline.

For 2D correlation analysis between ECoG and laminar probe signals, data was band pass filtered with a second order Butterworth filter (10-20 Hz or 100-200 Hz), z-scored and corrected for median channel activity per channel. The first second of the data after stimulus onset from all 8 stimulus orientation conditions was used to calculate the Pearson correlation coefficients for all ECoG channels against individual laminar probe channels.

To estimate the retinotopic alignment expected from random probe placement, we performed a Monte Carlo simulation based on the empirical RF location of the V1 laminar probe and the retinotopic map obtained from MT µECoG recordings. The median RF location of the V1 laminar probe was used as a reference. Random MT probe locations were generated by uniformly sampling RF coordinates from the MT µECoG retinotopic map. For each sample, the Euclidean distance to the V1 reference was calculated and this was repeated 10,000 times to obtain the expected distance distribution, against which the observed V1–MT RF distance was compared.

For analysis of spatiotemporal activity patterns (Fig. 3i-k) LFP signals were band-pass filtered in the gamma range (60–90 Hz) using a third-order Butterworth filter. The analytic signal was computed using the Hilbert transform and averaged across trials. Envelope signals were smoothed using a Gaussian kernel (σ = 250 ms). Signal-to-noise ratio (SNR) was calculated as the ratio of peak envelope amplitude to baseline activity (mean of first and last 50 ms). Three channels (Fig. 3i) with SNR < 1.5 were excluded and linearly interpolated. To track activity across time and space, data were thresholded at each time point (≥80% of normalized amplitude), and the center of mass of the largest region was computed using amplitude-weighted coordinates to obtain the trajectory of activation across the array.

## Acknowledgements

We thank Thomas Wunderle and Gustavo Rohenkohl for their assistance during a portion of the experiments. This work was supported by DFG (SPP 1665 FR2557/1-1, FOR 1847 FR2557/2-1, FR2557/5-1-CORNET, FR2557/6-1-NeuroTMR, FR2557/7-1-DualStreams to P.F.), EU (HEALTH-F2-2008-200728-BrainSynch, FP7-604102-HBP, FP7-600730-Magnetrodes to P.F.), National Institutes of Health (1U54MH091657-WU-Minn-Consortium-HCP to P.F.), the LOEWE program (NeFF to P.F.) and the University of Zurich (project K-41220-04, C.L.).

## Author contributions

P.J, P.F., and C.M.L. designed research; B.R., R.L. and T.S. fabricated the polyimide electrode arrays, P.J. and C.M.L. performed experiments; P.J. and C.M.L. analyzed data; P.F. and C.M.L. provided supervision, C.M.L. and P.J. wrote the paper with input from the other authors.

