## Supplemental notes and figures for "A hybrid micro-ECoG for functionally targeted multi-site and multi-scale investigation"

### Stability and Reusability of Hybrid µECoG Arrays (related to Figure 3–5)

The hybrid µECoG arrays were reused across multiple acute experiments that each lasted up to several days. Recordings from cat visual cortex spanned approximately 136.6 hours (~5.7 days) and recordings from marmoset visual cortex spanned approximately 82.6 hours (~3.4 days) under continuous general anesthesia. Data shown in Figure 5 (cat cortex) were obtained using the same array as in Figure 3 (marmoset cortex), approximately 8 months later. Between experiments, the arrays were cleaned and stored dry. Qualitatively, both experiments yielded clear visually driven responses and well-defined receptive fields, indicating that the µECoG arrays remained functional after being used.

Quantitative comparison between experiments might be confounded due to species differences and individual variability in brain anatomy, as well as differences in electrical reference quality and noise levels for each experiment. However, it is nevertheless possible to estimate changes in signal quality within each experiment by calculating RMS values of the baseline period (first 200 ms of each trial, before stimulus onset). To reduce the influence of changes in neural signals on this noise estimate, we calculated RMS values for the high-pass component of the LFP (>150 Hz), because most of the energy of physiological signals is contained in the lower frequency band (Buzsáki et al. 2012).

RMS levels remained within a similar order of magnitude across recording days (Fig. S6 a,b). Spearman correlation of the median RMS values over time indicated no significant increase across recording blocks (Cat: $\rho = 0.188$, $p = 0.608$; Marmoset: $\rho = 0.250$, $p = 0.595$). To further assess the stability of the array, we also computed the percentage of outlier channels per recording timepoint (defined as ±3 SD from the mean within each recording). The Spearman correlation again showed no significant trend (Cat: $\rho = 0.188$, $p = 0.604$; Marmoset: $\rho = 0.359$, $p = 0.438$). These results indicate that the arrays remained functional across days within experiments and across multiple experiments.

To exclude any remaining influence of physiological effects on the analysis of array quality and reusability, we also compared impedance values before the first usage with values after it had been used in 5 experiments over approximately eight months (Fig. S6 c). Median impedance increased by 13.8% from 202 kOhm to 230 kOhm (Wilcoxon rank-sum test, $p = 7.84 \times 10^{-8}$; mean increased from 204 kOhm to 279 kOhm). Only 16 of the 240 channels (6.7%) showed very high impedance values (>800 kOhm) after repeated usage. Accordingly, 224 of 240 channels (93.3%) appeared functionally intact based on this impedance criterion.

While the observations above indicate that the arrays remained functional across multiple days within experiments and after repeated use, chronic preparations would be needed for systematic evaluation of long-term recording stability and should be addressed in future work.

### Supplemental Figures

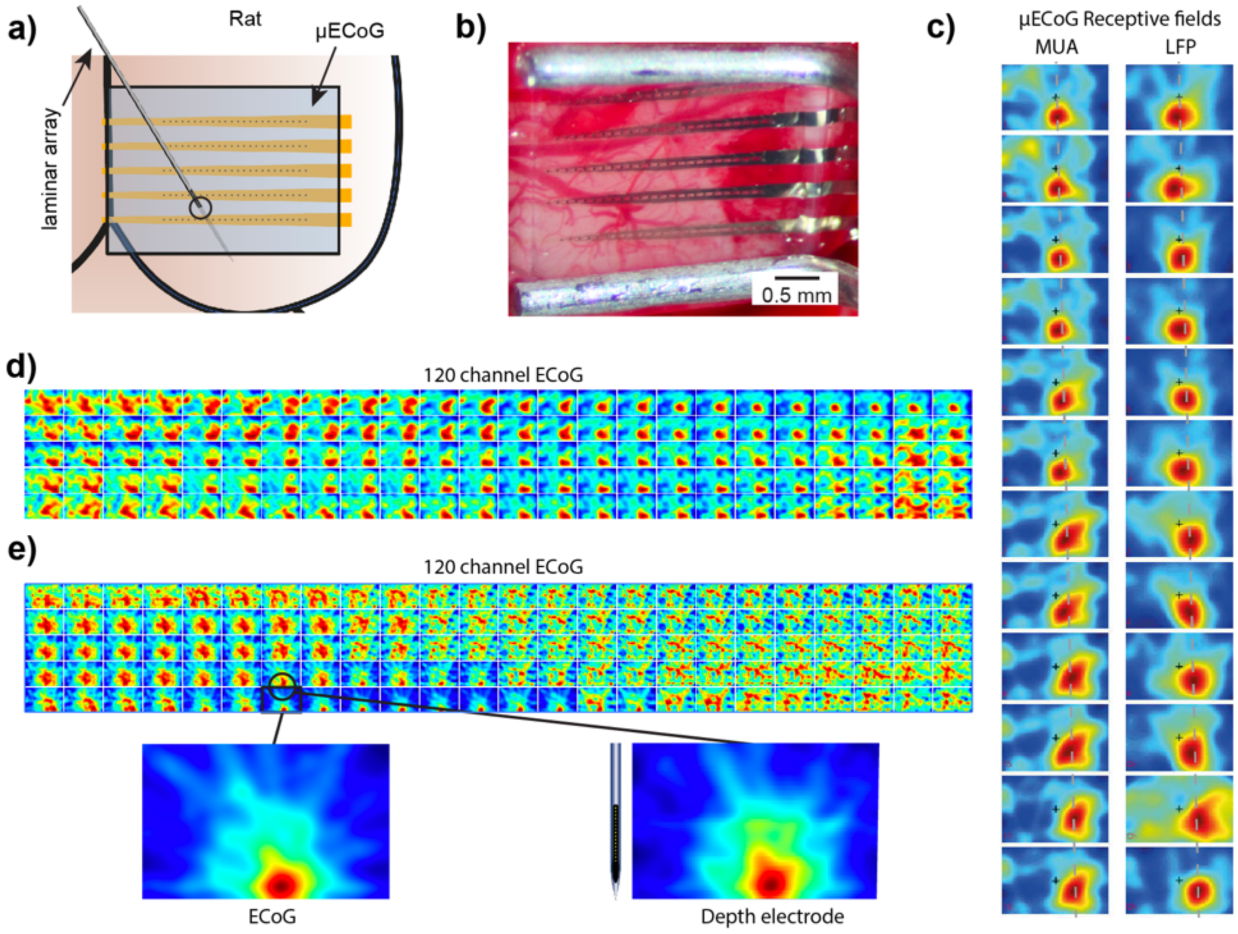


**Figure S1: Hybrid µECoG functional mapping of rodent visual cortex** (A) Illustration of the hybrid µECoG placed over the posterior dorsal cortex of the rat brain with a laminar array penetrating the µECoG into the cortical depth. b) Photograph of a hybrid µECoG in position above the rat posterior cortex. c) Comparison of RF estimates from MUA or LFP (power in the 60-100 Hz band) recorded from a µECoG array placed above the rat visual cortex. d) Example visual RF estimates across the rat visual cortex using a hybrid µECoG. e) Example of visual receptive fields estimated from µECoG recordings (left bottom panel) and from a laminar electrode array inserted through the µECoG to the cortical depth (right bottom panel).

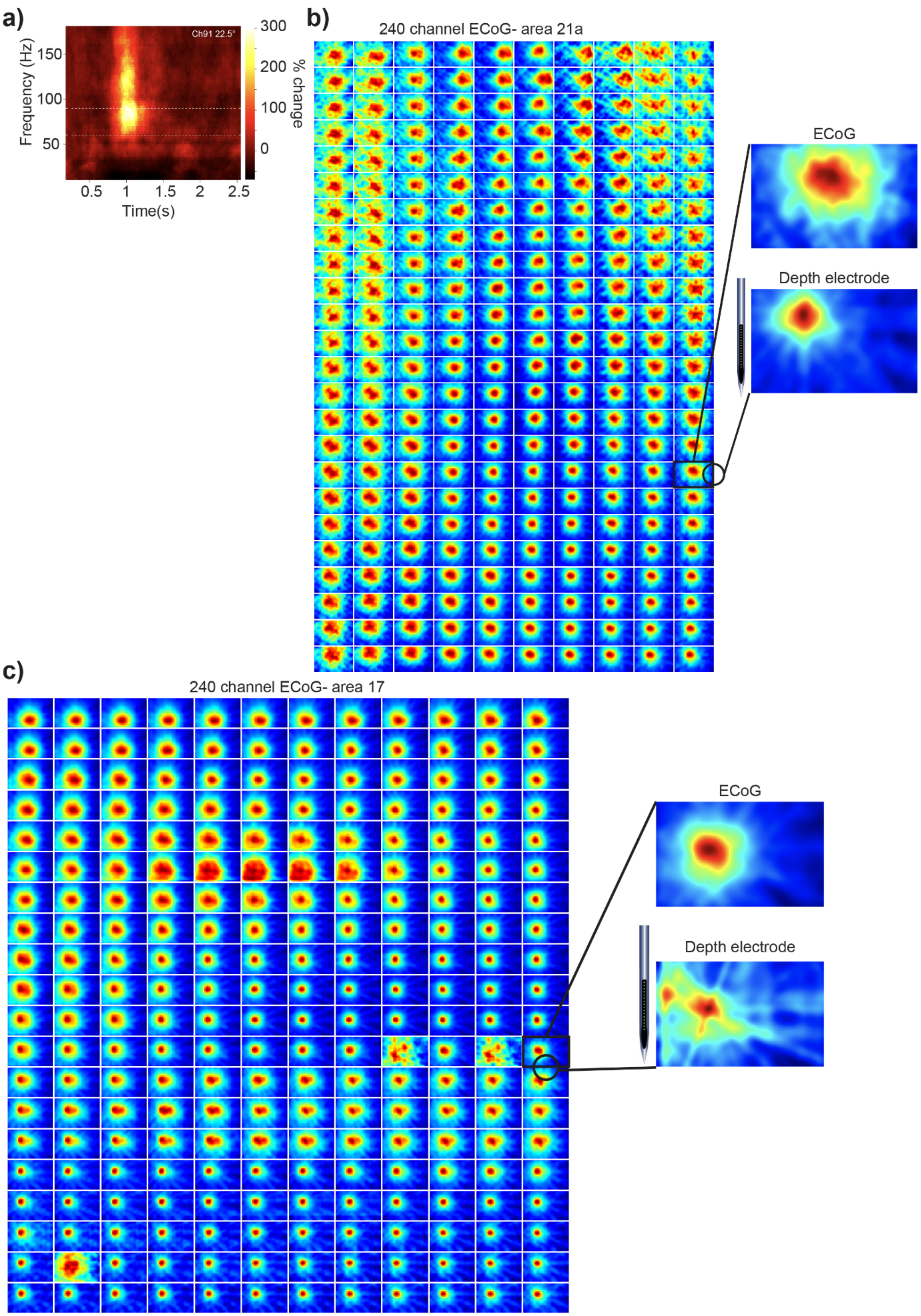

a)
Ch91 22.5°
Frequency (Hz)
150
100
50
0.5
1
1.5
2
2.5
Time(s)
300
200
100
0
% change
b)
240 channel ECoG- area 21a
ECoG
Depth electrode
c)
240 channel ECoG- area 17
ECoG
Depth electrode

**Figure S2: Receptive fields estimated from μECoG in areas 17/18 and 21a of the cat visual cortex. a)** Example time-frequency plot of the LFP response to visual stimulation with the RF mapping protocol. A transient increase in power in the frequencies above 60 Hz is visible. For the LFP RF mapping, we used the gamma-band power to estimate the RF for each μECoG electrode. **b)** The RF for each μECoG electrode for the array placed over area 17/18 are plotted as a function of position on the ECoG array. (right) The RFs estimated using LFP from the ECoG agree well with RFs estimated from neuronal MUA activity recorded in the depth using linear electrode arrays. **c)** RFs mapped from LFP power for the ECoG in area 21a. Subplots on the right show a magnified view of the RFs from the depth electrode (bottom), inserted into the cortex through the ECoG and the receptive field of the immediately neighboring EGoG electrode (top), revealing very similar spatial selectivity. Note that the RF maps for the area 17 and 21a ECoG arrays are drawn in scale in coordinates of the visual display and do not reflect the relative physical dimensions of the ECoG arrays.

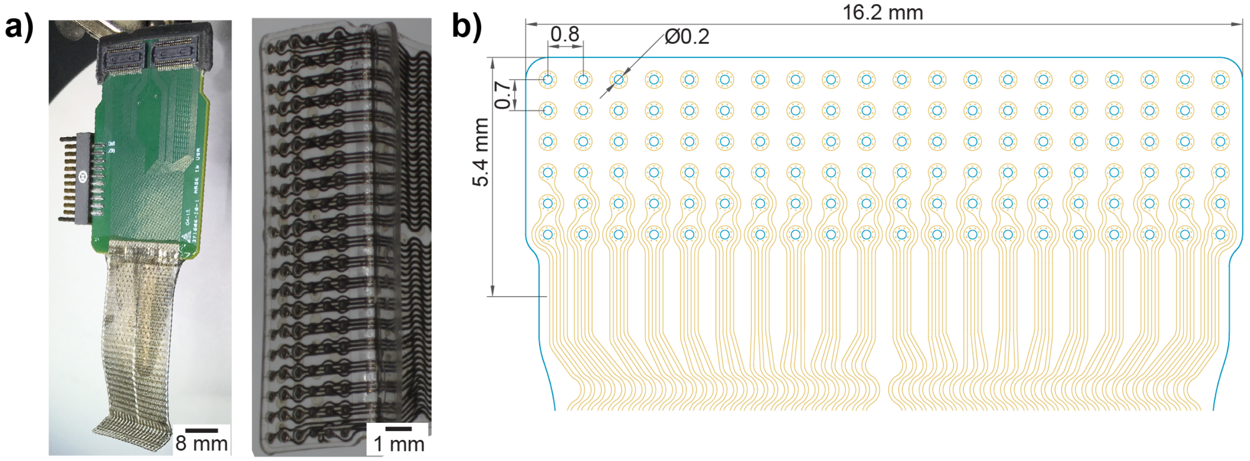


**Figure S3: Silicone μECoG fabricated by CorTec GmbH.** a) Left, Photograph of assembled μECoG and connector board. Right, Zoomed photograph showing the μECoG array electrodes and traces. Compared to our hybrid μECoG, more of the field of view is obscured by electrodes and wiring, because of the lower resolution of the pure PDMS μECoG. b) Schematic layout of the CorTec PDMS μECoG design.

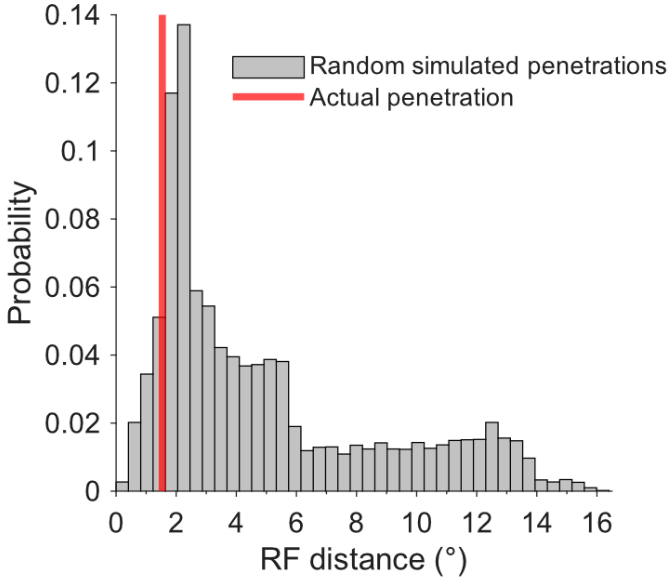


**Figure S4: Monte Carlo estimate of retinotopic alignment for MT probe placement**. Distribution of RF distances between the V1 laminar probe and simulated randomly sampled locations from the MT µECoG retinotopic map (grey histogram). The red line indicates the observed RF distance for the actual MT probe penetration during the experiment. The observed distance (1.54°) was smaller than 91.3% of distances obtained from random sampling.

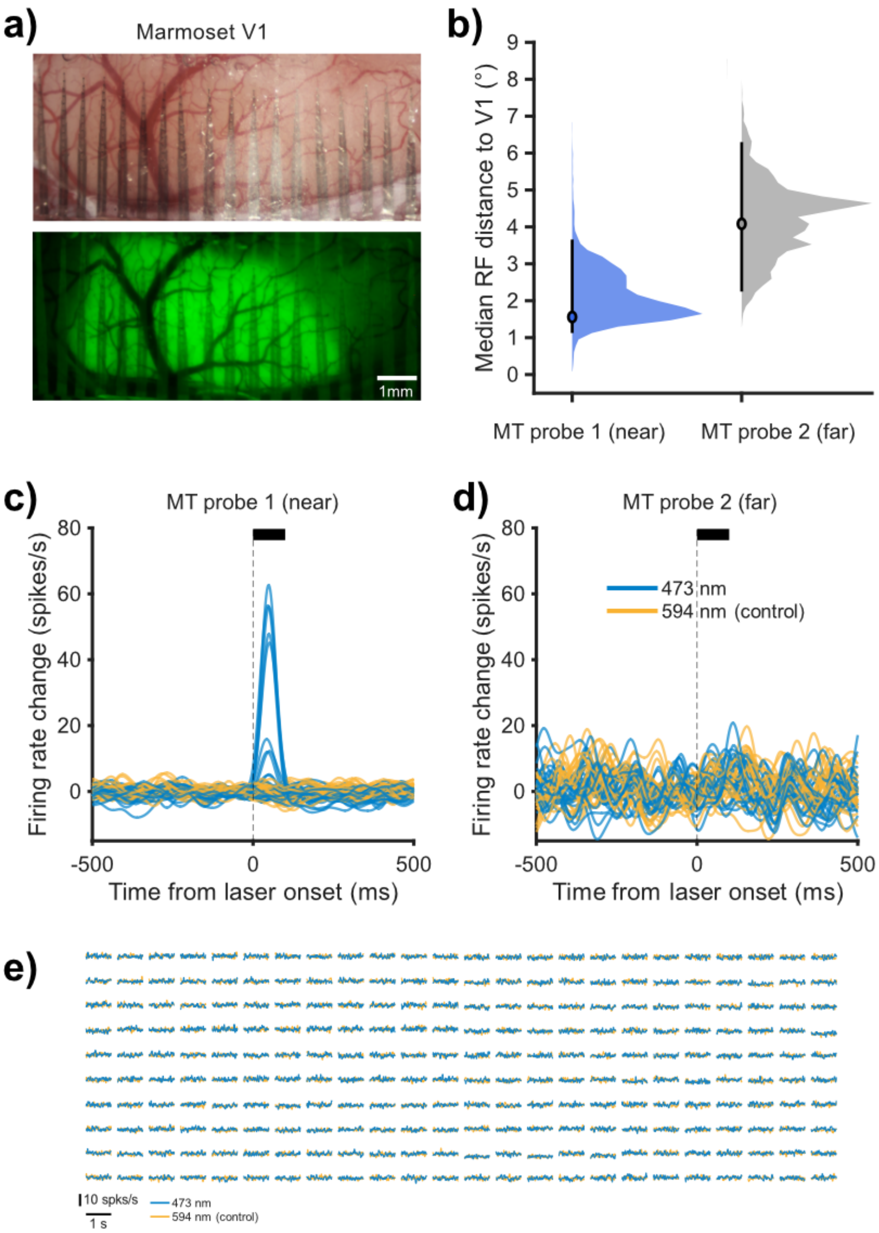


**Figure S5: Optogenetic responses in MT depend on retinotopic alignment.** **a)** Photograph of the hybrid μECoG over marmoset V1 (top), and epifluorescence image (bottom) showing the virally expressed fluorescence protein (eYFP**).** **b)** Bootstrap distributions of the median RF distance between the V1 laminar probe and two MT probes. The MT probe closer in retinotopic space shows a smaller RF distance. Points indicate observed medians; error bars indicate 95% bootstrap confidence intervals. **c)** Optogenetically evoked spiking responses for the MT probe with smaller RF distance to V1 ("near") with stimulation (blue, 473 nm) and control condition (yellow, 594 nm). **d)** Responses for the MT probe with larger RF distance ("far"), showing no optogenetically evoked activity.**e)** Absence of multiunit (MUA) responses on the μEcoG in area MT to optogenetic stimulation in V1.

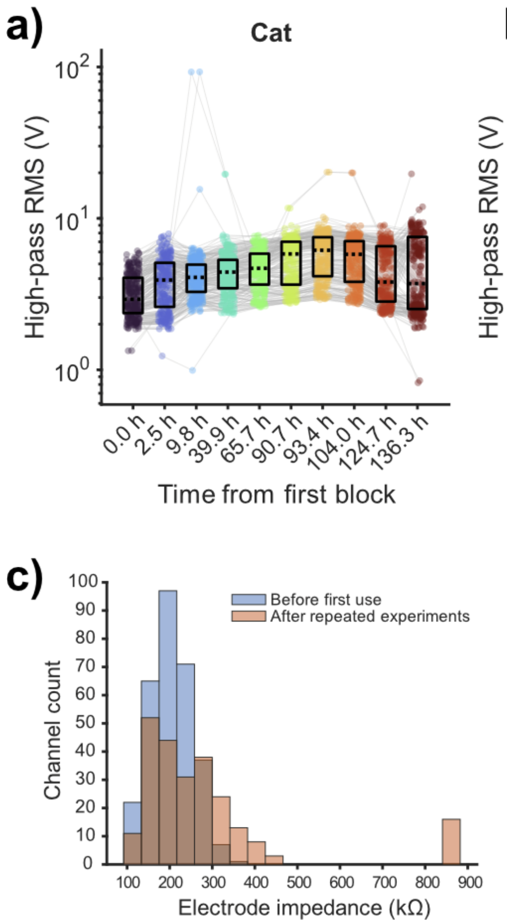


**Figure S6: Signal quality across recording days and impedance values changes. a)** High-pass RMS values (>150 Hz) of baseline activity (first 200 ms before stimulus onset) across recording blocks in cat cortex. Each dot represents one channel, grey lines connect repeated measurements from the same channel across time, and boxplots summarize the distribution at each time point (median and interquartile range).**b)** Same analysis for marmoset cortex. RMS values remained within a similar range across recording days, and no significant correlation with time was observed (cat: $\rho = 0.188$, $p = 0.608$; marmoset: $\rho = 0.250$, $p = 0.595$). **c)** Distribution of electrode impedance values before first use and after repeated use across five experiments (~8 months). Median impedance increased (202 kΩ to 230 kΩ; Wilcoxon rank-sum test, $p = 7.84 \times 10^{-8}$). 16 out of 240 channels (6.7%) showed high impedance values (>800 kΩ) after repeated use.

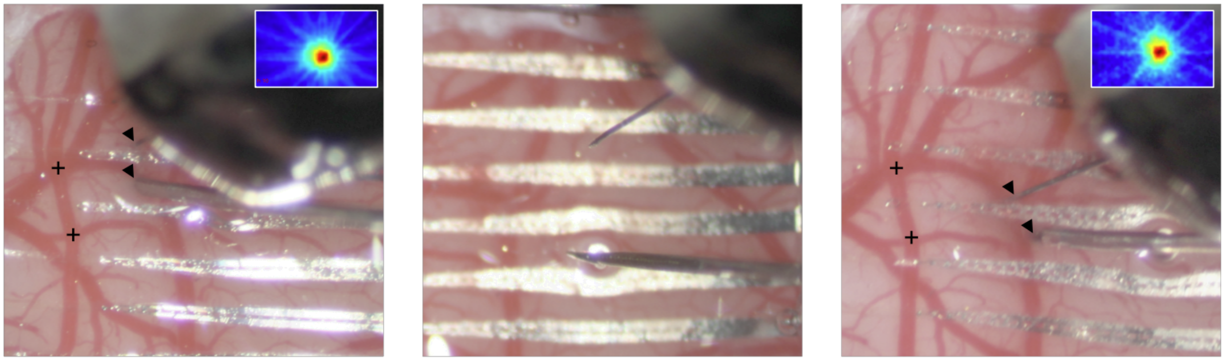

**Figure S7. Repeated penetration of laminar probes through the hybrid μECoG.** The soft PDMS encapsulation of the hybrid μECoG permits repeated penetration by depth electrodes, resealing after probe retraction. **Left:** Two laminar probes inserted through the μECoG in marmoset area MT. The upper probe is a 32-channel silicon probe (Atlas Neurotechnology), and the lower probe is a 24-channel Plexon V-probe. Insertion sites are indicated by arrowheads. **Center:** The Atlas probe is fully retracted while the V-probe remains minimally inserted. **Right:** Both probes are reinserted through the μECoG. Insets (left and right) show example receptive fields from the same channel of the 32-channel Atlas probe before and after re-insertion. Black crosses in the left and right panels indicate landmarks of corresponding positions on the cortical surface.
